# SMART: Statistical Mitogenome Assembly with Repeats

**DOI:** 10.1101/795633

**Authors:** Fahad Alqahtani, Ion I. Măndoiu

**Affiliations:** Computer Science & Engineering Department, University of Connecticut, Storrs, CT, USA; National Center for Arteficial Intelligence and Big Data Technology, King Abdulaziz City for Science and Technology, Riyadh, Saudi Arabia

**Keywords:** *De novo* assembly, mitogenomes, repeats

## Abstract

By using next-generation sequencing technologies it is possible to quickly and inexpensively generate large numbers of relatively short reads from both the nuclear and mitochondrial DNA contained in a biological sample. Unfortunately, assembling such *whole-genome sequencing (WGS)* data with standard *de novo* assemblers often fails to generate high quality mitochondrial genome sequences due to the large difference in copy number (and hence sequencing depth) between the mitochondrial and nuclear genomes. Assembly of complete mitochondrial genome sequences is further complicated by the fact that many *de novo* assemblers are not designed for circular genomes, and by the presence of repeats in the mitochondrial genomes of some species.

In this paper we describe the *Statistical Mitogenome Assembly with Repeats (SMART)* pipeline for automated assembly of complete circular mitochondrial genomes from WGS data. SMART uses an efficient coverage-based filter to first select a subset of reads enriched in mtDNA sequences. Contigs produced by an initial assembly step are filtered using BLAST searches against a comprehensive mitochondrial genome database, and used as “baits” for an alignment-based filter that produces the set of reads used in a second *de novo* assembly and scaffolding step. In the presence of repeats, the possible paths through the assembly graph are evaluated using a maximum-likelihood model. Additionally, the assembly process is repeated a user-specified number of times on re-sampled subsets of reads to select for annotation the reconstructed sequences with highest bootstrap support.

Experiments on WGS datasets from a variety of species show that the SMART pipeline produces complete circular mitochondrial genome sequences with a higher success rate than current state-of-the art tools, even from low coverage WGS data. The pipeline is available through an easy-to-use web interface at https://neo.engr.uconn.edu/?tool_id=SMART.

## Background

The mitochondria are cellular organelles often called powerhouses of the cell due to their key role in the production of adenosine triphosphate (ATP). Found in most eukaryotic organisms, the mitochondria have their own circular genomes. They are inherited maternally in most animals, and are typically present in thousands of copies in the cytoplasm of each cell, although the copy number varies between cells of different tissues [1]. Single nucleotide polymorphisms in mitochondrial genomes have long been used for tracking human migrations [2]. Mitochondrial DNA (mtDNA) mutations and *heteroplasmy* (simultaneous presence of multiple mitochondrial sequences in a cell) have also been associated with human diseases [3]. Moreover, mtDNA analysis can be a useful tool in forensics, especially when a crime scene sample contains degraded DNA not suitable for nuclear DNA tests [4]. Finally, mitochondrial genome sequences can be used for evolutionary studies of non-model species for which nuclear genomes are not yet available [5].

Mitochondrial DNA can be experimentally separated from the nuclear DNA and sequenced independently but such protocols are laborious [6]. More commonly, the mitogenomes are assembled from *Whole Genome Sequencing (WGS)* data, which consists of reads generated from both the nuclear and mitochondrial genomes [6]. Unfortunately, assembling WGS data with standard *de novo* assemblers often fails to generate high quality mitochondrial genome sequences due to the large difference in copy number (and hence sequencing depth) between the mitochondrial and nuclear genomes [7]. This has led to the development of specialized tools for reconstructing mitochondrial genomes from WGS data. Although assembly of mitochondrial genomes from long-read WGS data has been demonstrated [8], the high coverage required (> 50×) and the relatively high cost of long-read sequencing make this approach uncommon.

Consequently, most of the existing tools for mtDNA assembly have focused on the most abundant type of WGS data currently available, which consists of relatively short reads, typically around 100bp. These tools can be grouped into three main categories: reference-based, seed-and-extend, and *de novo* assembly. Reference-based methods such as MToolBox [9] require the mtDNA sequence of the species of interest or a closely related species. These approaches have the lowest running time and memory requirements, but cannot be used for non-model organisms for which such a reference is not available. MITObim [7] and NOVOPlasty [10] are two tools that implement the seed-and-extend approach for reconstructing circular organelle genomes including mitogenomes (MITObim also implements reference-based assembly). The results in [7] show that the seed-and-extend approach can successfully assemble mitochondrial genome sequences starting from a very short seed such as the sequence of the Cytochrome c oxidase subunit I (COI) gene, which is commonly used as a DNA barcode for animals [11] and is widely available for numerous species [12]. However, due to their inherently greedy approach, seed-and-extend methods have difficulty handling repetitive regions present in some mitochondrial genomes [5]. Norgal [6] is a recent tool implementing a *de novo* approach to mitochondrial genome reconstruction from WGS data, without the need for either a reference or a seed sequence. A similar approach is used by plasmidSPAdes [13] for *de novo* assembly of circular plasmid genomes from WGS data. Although these tools are broadly applicable, they can have prohibitive running times and may still fail to reconstruct complete mitogenomes, particularly in the presence of repeats shared between the nuclear and organelle genomes [14].

In this paper we describe the *Statistical Mitogenome Assembly with Repeats (SMART)* pipeline for *de novo* assembly of complete circular mitochondrial genome sequences from WGS data. To ensure a high assembly success rate even from low-coverage WGS data and in the presence of repeats, SMART employs several novel techniques. First, SMART uses an initial coverage-based filtering step to enrich for mtDNA reads. Although similar filtering steps are included in other *de novo* pipelines such as Norgal [6] and plasmidSPAdes [13], the approach taken in SMART is different. Norgal and plasmidSPAdes attempt to remove reads from the nuclear genome by performing an assembly of the full set of reads and then using the read coverage of the longest contigs to estimate the coverage of the nuclear genome. On the other hand, SMART estimates the mean and standard deviation of mtDNA k-mer counts in WGS reads based on a seed sequence, then positively selects reads with observed k-mer counts falling within three standard deviations of the estimated mean. As shown in the Results section, the positive selection approach of SMART is robust to large variations in mtDNA read content and yields higher enrichment for mtDNA reads than the negative selection implemented by Norgal for low-depth WGS datasets. Furthermore, positive selection based on k-mer counting removes the time consuming assembly of all WGS reads required by Norgal and plasmidSPAdes. Second, SMART iteratively refines the set of selected reads and uses a maximum likelihood model to increase assembly accuracy. Reads passing the coverage-based filter are assembled using Velvet to generate a preliminary set of contigs. Preliminary contigs are themselves filtered using BLAST searches against a comprehensive mitochondrial genome database to extract likely mitochondrial contigs, which are then used as “baits” for an alignment-based filter that produces a refined set of reads used in a second *de novo* assembly and scaffolding step using SPAdes [15] and SSPACE [16]. This process is repeated if the assembly graph is not Eulerian, and, in the presence of repeats, the possible paths through the assembly graph are evaluated using the ALE maximum-likelihood model [17]. Finally, the assembly process is repeated a user-specified number of times on re-sampled subsets of reads to select for annotation the assembled sequences with highest bootstrap support.

## Methods

### SMART pipeline interface

The SMART pipeline is deployed on a customized instance of the Galaxy framework [18] and can be accessed at https://neo.engr.uconn.edu/?tool_id=SMART. The pipeline was designed for processing paired-end WGS reads in fastq format. In addition to two fastq files, the user specifies the sample name and a seed sequence in fasta format (see Fig. 1 for a screenshot of the web interface). Optional parameters include the number of bootstrap samples (default 1), the number of read pairs per bootstrap sample (default 10M), the *k*-mer size (default 31), the number of CPU threads (default 16), and the genetic code to be used for annotation (default: vertebrate mitochondrial code). All experimental results in next section use the default setting unless otherwise noted.

**Figure 1:**
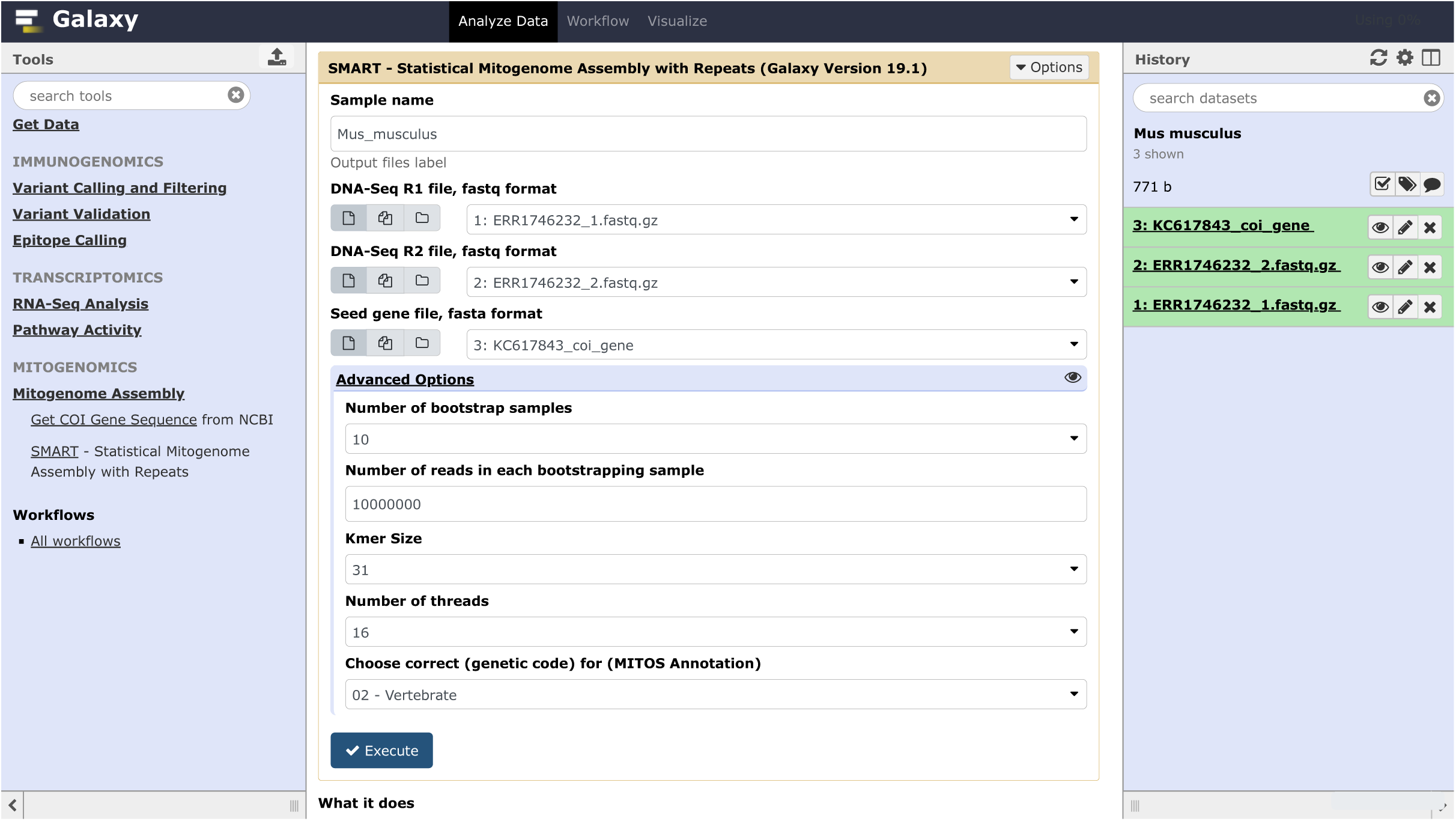
Web-based interface for the SMART pipeline, publicly accessible at https://neo.engr.uconn.edu/?tool_id=SMART.

Upon successful completion SMART generates three files:

- A zip file including the consensus sequence for each cluster in fasta and GenBank formats as well as MITOS [19] annotation files.
- A detailed pdf report that includes statistics and visualizations of various pipeline steps and the final mitogenome annotations. Sample reports from the 10 non-human species datasets discussed under Results are included in Additional File 1.
- A detailed log file that contains additional information including timing for each pipeline step.

### Seed selection

Similar to seed-and-extend tools such as MITObim [7] and NOVOPlasty [10], SMART requires as input a seed sequence. However, unlike MITObim and NOVO-Plasty, SMART uses the seed sequence only for estimating mtDNA *k*-mer coverage and implementing an efficient coverage-based read filter – all assembly steps are performed *de novo* using de Bruijn graph assemblers Velvet [20] and SPAdes [15]. As shown in the Results section, high quality mitogenome sequences can be obtained using seed sequences as short as a few hundred bases. Additionally, although seed sequences from the same species are preferable, assembly can succeed even with seeds from closely related species. For ease of use SMART includes a tool for importing to Galaxy seed sequences from GeneBank based on their accession number. A widely available seed sequence is the Cytochrome c Oxidase I (COI) mitochondrial gene, which is commonly employed as a DNA barcode for species identification [21]. The largest repository of COI sequences is the Barcode of Life Data System (BOLD) [12], which currently includes more than 1,419,768 public barcode sequences from 118,358 animal species.

### SMART workflow

The main stages of the SMART pipeline (see Fig. 2) are as follows:

**Figure 2:**
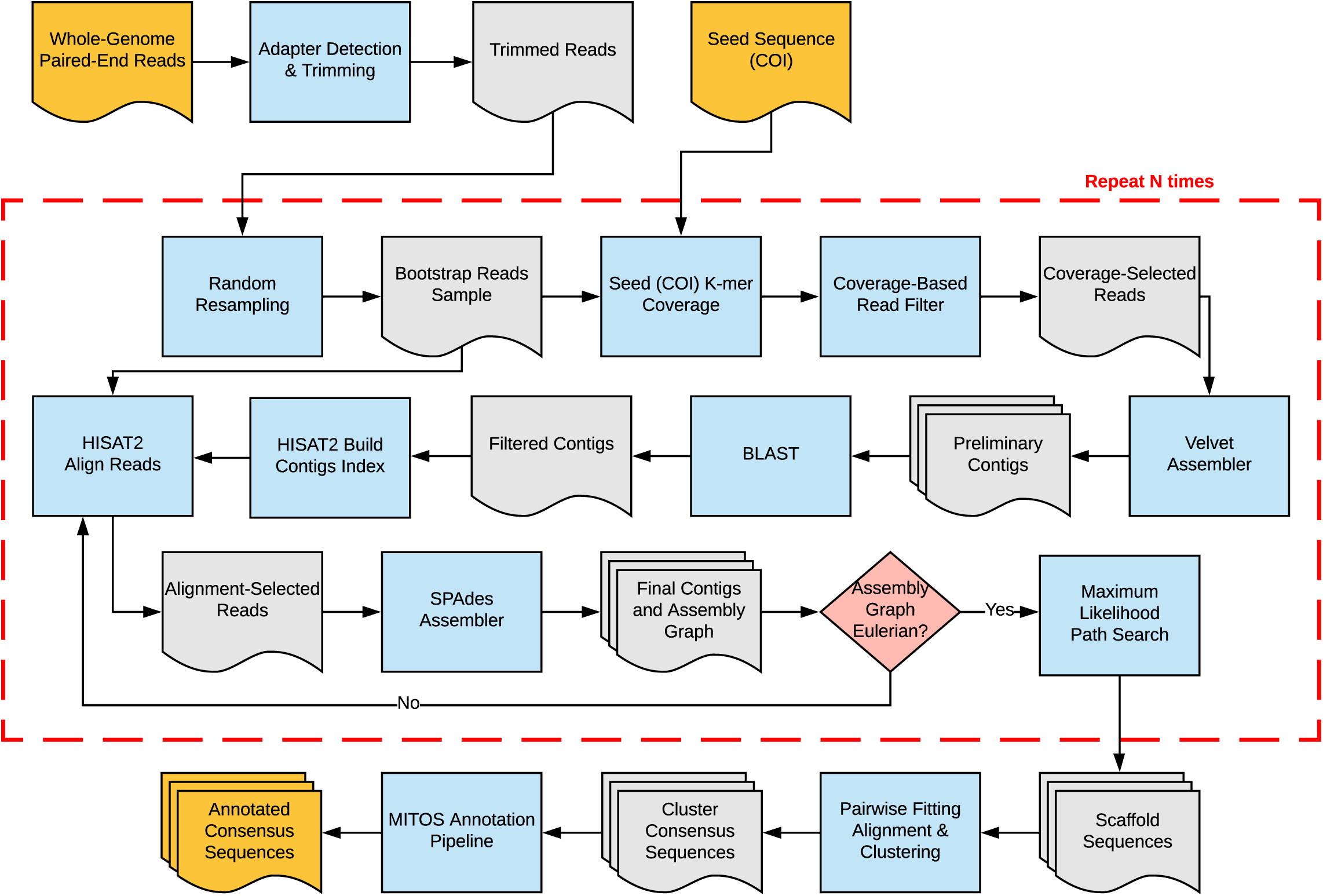
SMART pipeline flowchart.

1. Automatic adapter detection and trimming.
2. Random resampling of a user-specified number of trimmed read pairs.
3. Coverage-based filtering of the reads based on the seed sequence.
4. Preliminary assembly of reads passing the coverage filter.
5. Filtering of preliminary contigs by BLAST searches against a local mitochondrial database.
6. Secondary read filtering by alignment to preliminary contigs that have significant BLAST matches.
7. Secondary *de novo* assembly.
8. Iterative scaffolding and gap filling based on maximum likelihood.
9. Prediction and annotation of mitochondrial genes.

By default the above process is executed only once, but SMART users can specify the number of times steps 2-8 should be repeated to compute the bootstrap support for the assembled sequences. When the number of bootstrap samples is greater than one, the resulting circular sequences are clustered based on their pair-wise distances, and the annotation step is performed on the consensus sequence obtained for each cluster. Details on each of the workflow steps are provided below.

#### Adapter detection and trimming

Due to variability in sample quality and library preparation protocols, next-generation sequencing data can include substantial amounts of sequencing errors and other technical artifacts such as adapter contamination. Since such artifacts can negatively impact the quality of downstream analyses including *de novo* assembly, there are numerous tools that can be used for quality checking and filtering WGS data [22, 23]. However, many of these tools require iterative user intervention [22, 23]. To minimize user involvement, in SMART we have only incorporated automatic adapter detection and removal. Specifically, we detect and trim adapters using tools included in the IRFinder package [24]. These tools take advantage of the fact that for paired-end WGS data adapter sequences are included in both reads when the target DNA fragment is shorter than the read length. This allows both highly accurate automatic detection of adaptor sequences from a small data sample and very precise (single base resolution) adapter trimming.

#### Random read re-sampling

After adapter trimming, SMART generates a user-specified number of bootstrap samples by re-sampling. These samples are generated using the FASTQ-SAMPLE tool from the FASTQ-TOOLS package [25].

#### Coverage-based read filtering

The aim of this step is to filter out nuclear reads by taking advantage of the difference in copy number between the nuclear and mitochondrial genomes. Due to this difference, the counts of *k*-mers that originate from the mitocondrial genome are expected to be much higher than that of *k*-mers from the nuclear genome, with the possible exception of *k*-mers that originate from nuclear genome repeats with similar copy number. To implement a filter based on this observation, we use the Jellyfish package [26] to efficiently count the number of times each *k*-mer appears in the reads of the bootstrap sample. To account for sequencing errors and low degrees of dissimilarity between the sequenced mitogenome and the seed sequence, for each *k*-mer of the seed sequence we augment the observed Jellyfish count by adding the counts of the *k*-mers at Hamming distance one. Although most seed sequence *k*-mers are expected to have high augmented counts, *k*-mers from regions of the seed sequence that have high dissimilarity to the homologous region of the sequenced mitogenome will still have zero or near-zero augmented counts. Consequently, we use the MCLUST package [27] to fit a two-component Gaussian mixture model to the one-dimensional distribution of augmented *k*-mer counts, and use the mean *µ* and standard deviation *σ* of the upper component as the estimate for the corresponding mtDNA *k*-mer count statistics.

To efficiently extract putative mitochondrial reads, a hash table is populated with *all* read *k*-mers (not just seed sequence *k*-mers) that have a count within 3 standard deviations of the estimated mtDNA *k*-mer count mean, i.e., all *k*-mers *x* for which

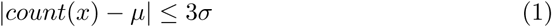

A read of length *l* in the bootstrap sample is then considered to be of mitochondrial origin if at least *l* − (2*k* − 1) of its *k*-mers are found in the hash table, i.e., satisfy (1). We allow up to 2*k* − 1 of the *k*-mers to violate (1) to ensure that we retain mitochondrial reads with a single sequencing error, since such an error can create up to 2*k* − 1 “novel” *k*-mers that would not match the expected count distribution. Both reads in a pair must satisfy this test in order for the pair to be kept; if either one of the reads or both fail the test the pair is removed. Experimental results in the next section show that the coverage-based filter typically leads to a substantial enrichment in mitochondrial reads, even when the coverage estimates are based on relatively few reads and short seed sequences.

#### Preliminary assembly

The goal of this step is to generate longer contigs from the enriched set of mitochondrial reads that pass the coverage-based filter. For time and memory usage efficiency, we use Velvet [20], a fast short read assembler based on de Bruijn graphs. Since some nuclear genome reads are expected to pass the coverage-based filter, the output of Velvet is typically a mixture of mitochondrial and nuclear genome contigs.

#### Preliminary contig filtering

The aim of this step to filter out nuclear genome contigs. This is accomplished by searching each contig against a local database of 8,376 complete eukaryotic mitogenomes downloaded from NCBI by using nucleotide-nucleotide BLAST 2.7.1+. As result of this step we retain only contigs that have hits with an E-value of 10^−10^ or less.

#### Alignment-based read filtering

The aim of this step to pull out additional mitochondrial reads that are missed by coverage filter using an alignment-based approach reminiscent of the seed-and-extend approaches. To implement this step efficiently, we build an index for the contigs with significant BLAST matches, then align all bootstrap reads against the index by using HISAT2, a fast and sensitive aligner for NGS reads [28]. Since the set of preliminary contigs is likely incomplete, both reads in a pair are kept if one of them is aligned. Specifically, all reads in a bootstrap sample are aligned using HISAT2 as single reads, and the union of all read IDs is given to the seqtk tool [29] to pull from the bootstrap sample the read pairs that have at least one of the reads aligned.

#### Secondary assembly

The goal of this step to assemble a high-quality mitochondrial sequence using the reads that pass the alignment-based filtering. SMART performs the secondary assembly using SPAdes, a multi-kmer de Bruijn graph assembler with robust performance even in the presence of non-uniformities in read coverage [15].

Since mitochondrial genomes are circular, SMART checks if the results of SPAdes assembly is an Eulearian graph using a custom python script. If so, SMART moves to the scaffolding step, otherwise SMART repeats the alignment-based read filtering and secondary assembly using SPAdes for up to 5 iterations. As a result of these iterations, the number of selected reads and the length of the assembled contigs typically increase.

#### Scaffolding

When SPAdes produces an Eulerian assembly graph or the maximum number of iterations is reached, SMART begins the scaffolding step. SMART generates all paths of the Eulerian graph using a depth-first approach. For each explored path SMART generates a scaffold sequence by trying to overlap adjacent contigs or closing gaps between contigs using SSPACE [16]. To select the most likely assembly SMART uses the ALE tool [17] to compute the likelihood of each scaffold sequence and outputs the sequence with maximum likelhood. The ALE likelihood model is based on four sub-scores: placement scoring takes into account how well read sequences agree with the scaffold sequence, insert scoring assesses how well insert lengths implied by the alignments of paired reads match the expected insert length distribution, depth scoring reflects how well the read depth at each location agrees with the depth expected after GC-bias correction, and k-mer scoring shows how well k-mer counts of each contig match the multinomial distribution estimated from the entire assembly. The ALE likelihood assessment is particularly useful for selecting high-confidence assemblies when the Eulerian graph has duplicated contigs.

#### Clustering

When the users choose to use multiple bootstrap samples SMART must consolidate the results of different runs and compute the bootstrap support for the final set of sequences. The output of each run is either a circular scaffold sequence or a linear scaffold sequence in case the assembly graph is not Eulerian. Furthermore, the scaffold sequences produced in each run may be generated from either the forward or reverse strands. The first step in the SMART consolidation process is to compute for each pair of scaffold sequences an alignment score that accommodates any combination of circular and linear sequences and is invariant to strand choice and rotations of the circular sequences when present. We do so by using dynamic programming to compute the optimal fitting alignment under the edit distance scores between a duplicated version of the longest sequence (arbitrarily linearized in case it is circular) and both the shortest sequence (again arbitrarily linearized in case it is circular) and its Watson-Crick complement. The smaller of the two edit distances, which is computed in time proportional to the product between the lengths of the two sequences a pairwise score, has the desired invariance properties. Indeed, the duplicated sequence contains as substrings all possible linearizations of the longest string and the fitting alignment algorithm finds the substring that has minimum edit distance to the (arbitrarily selected) linearization of the second string. By computing the fitting alignment against both the shortest string and its Watson-Crick complement we ensure that one of the computations has the two strings in compatible orientations.

Once all pairwise distances are computed SMART runs the hierarchical clustering algorithm implemented by the hclust R package [30] on the edit distance matrix and automatically cuts the resulting dendogram into clusters based on silhouette scores. Sequences within each cluster are flipped to the same strand and rotated to a consistent linearization using MARS [31]. Finally, SMART runs the MAFFT multiple sequences alignment tool [32] to generate a consensus sequence for each cluster.

#### Annotation

Each cluster consensus sequence is annotated using the MITOS *de novo* mitochondrial genome annotation pipeline [19], which identifies protein coding genes based on BLAST searches against previously annotated protein sequences and annotates tRNA and rRNA genes based on manually curated covariance models capturing both sequence and secondary structure similarity to known sequences.

## Results and discussion

### Datasets

To assess the effectiveness of SMART and compare it with prior methods we used two groups of datasets. The first group, comprised of 8 human datasets, was used for a detailed assessment, including evaluation of the accuracy of various read filtering strategies and comparison with previous methods. Accession numbers and basic statistics for the human datasets are provided in Table 1. Six of the human datasets were generated using the WGS strategy, while the other two datasets were generated using Whole Exome Sequencing (WES), a sequencing protocol that has been previously shown to yield sufficient reads for mitogenome reconstruction using a reference based approach [33].

**Table 1:**
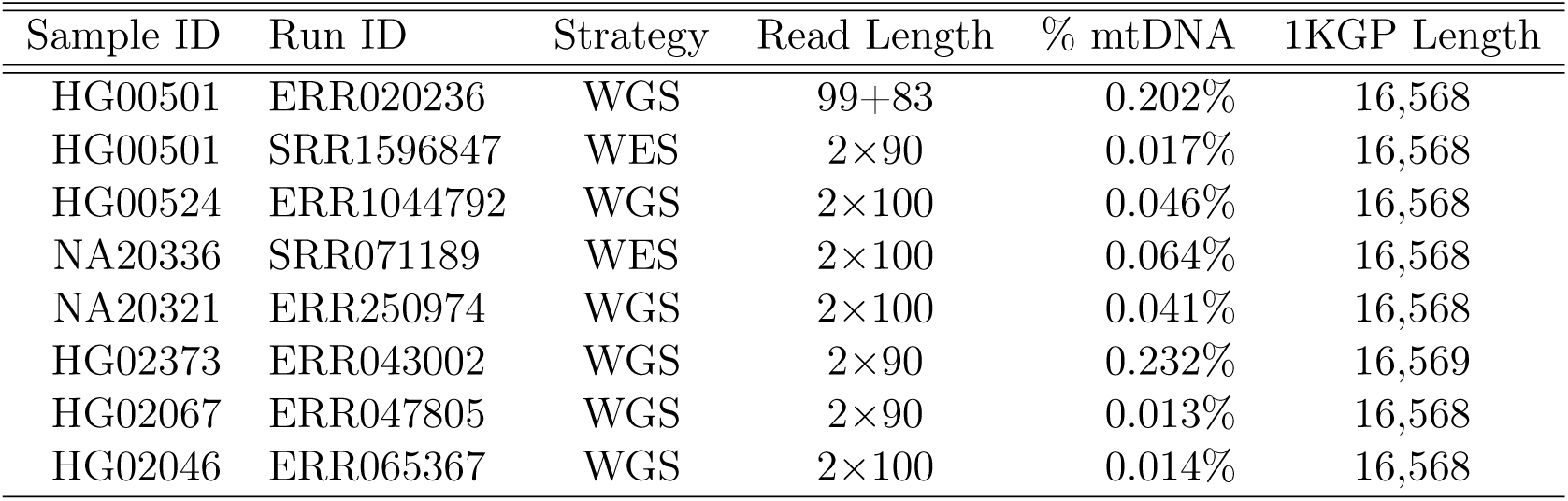
Human WGS and WES datasets used in evaluation experiments. All SMART runs were performed using between 2.5M and 25M read pairs and, unless otherwise noted, the 386bp-long human COI sequence with GenBank accession KC750830 as seed. mtDNA content was assessed by aligning the reads against the mtDNA sequence published for each sample by 1KGP.

The second group (Table 2) consists of WGS datasets from ten non-human species spanning the tree of life, including a primate dataset (*Pan troglodytes*), three other mammals (*Canis lupus, Capra hircus*, and *Mus Musculus*), a bird (*Grus Japonensis*), two frogs (*Rana temporaria* and *Xenopus laevis*), an insect (*Phlebotomus papatasi*), a plant (*Saccharina japonica*), and a fungus (*Aspergillus niger*).

**Table 2:** Non-human WGS datasets used in evaluation experiments, including the number of read pairs and accession numbers of COI sequences used as seeds for the SMART runs and the reference used for assessing assembly accuracy.

| Species | Run ID | Length | mtDNA | #Read Pairs | Seed | Reference |
| --- | --- | --- | --- | --- | --- | --- |
| *Aspergillus niger* | SRR1801279 | 2×150 | 4.258% | 100,000 | EF180096 | NC_007445 |
| *Canis lupus* | ERR690331 | 2×90 | 0.060% | 5,000,000 | KC985188 | KU644662 |
| *Capra hircus* | ERR2309151 | 2×90 | 0.035% | 10,000,000 | JQ735457 | MK341077 |
| *Grus japonensis* | SRR5992802 | 2×100 | 0.005% | 50,000,000 | KF939577 | FJ769847 |
| *Mus Musculus* | ERR1746232 | 2×100 | 0.653% | 10,000,000 | KC617843 | KY018919 |
| *Pan troglodytes* | ERR1709948 | 2×100 | 0.014% | 10,000,000 | AY544154 | KU308540 |
| *Phlebotomus papatasi* | SRR1997462 | 2×100 | 0.446% | 1,000,000 | MH780862 | NC_028042 |
| *Rana temporaria* | SRR2226373 | 2×101 | 0.068% | 5,000,000 | MF624326 | NC_042226 |
| *Saccharina japonica* | SRR2043182 | 2×101 | 0.141% | 3,000,000 | KC491236 | NC_040854 |
| *Xenopus laevis* | SRR3210975 | 2×150 | 0.005% | 40,000,000 | GQ862287 | HM991335 |

Since the human datasets originate from individuals sequenced as part of the 1000 Genomes Project (1KGP), the mtDNA sequences reconstructed by 1KGP were used as ground truth for assessing accuracy. For non-human datasets we assessed accuracy by using as ground truth the published mtDNA reference sequences for the respective species (accession numbers provided in last column of Table 2), which provide close (albeit not exact) approximations for the mitogenome sequences of the specific specimens used to generate the sequencing data.

A notable feature of the considered datasets is the highly variable percentage of reads of mitochondrial origin, ranging from 0.005% in *Grus Japonensis* to over 4% in *Aspergillus niger*. This percentage was estimated by aligning 25M read pairs (or the entire dataset if comprised of fewer than 25M read pairs) to the respective ground truth mtDNA sequence using bowtie2 [34]. The variability is not entirely a species effect – indeed, differences of more than an order of magnitude can be observed in the percentage of mtDNA reads in the human datasets. Most likely, other important contributing factors to this variability include DNA extraction and library preparation protocols [35], the tissue of origin [1], and the developmental stage of the sample [36].

### Read filtering accuracy

Figure 3 compares the accuracy of the read filter employed by Norgal with that of the coverage- and alignment-based filters of SMART on the human datasets described in Table 1. For each sequencing run, we varied the number of reads between 2.5M and 25M. For 2.5M read datasets, Norgal’s filter fails to select *any* mitochondrial reads on all but one of the runs. Although the Norgal filter’s performance improves somewhat at higher sequencing depth, with 3 out of the 8 runs achieving a non-zero True Positive Rate (TPR) for 25M reads, its Positive Predictive Value (PPV) remains close to zero, showing that the vast majority of reads that pass the Norgal filter have nuclear origin.

**Figure 3:**
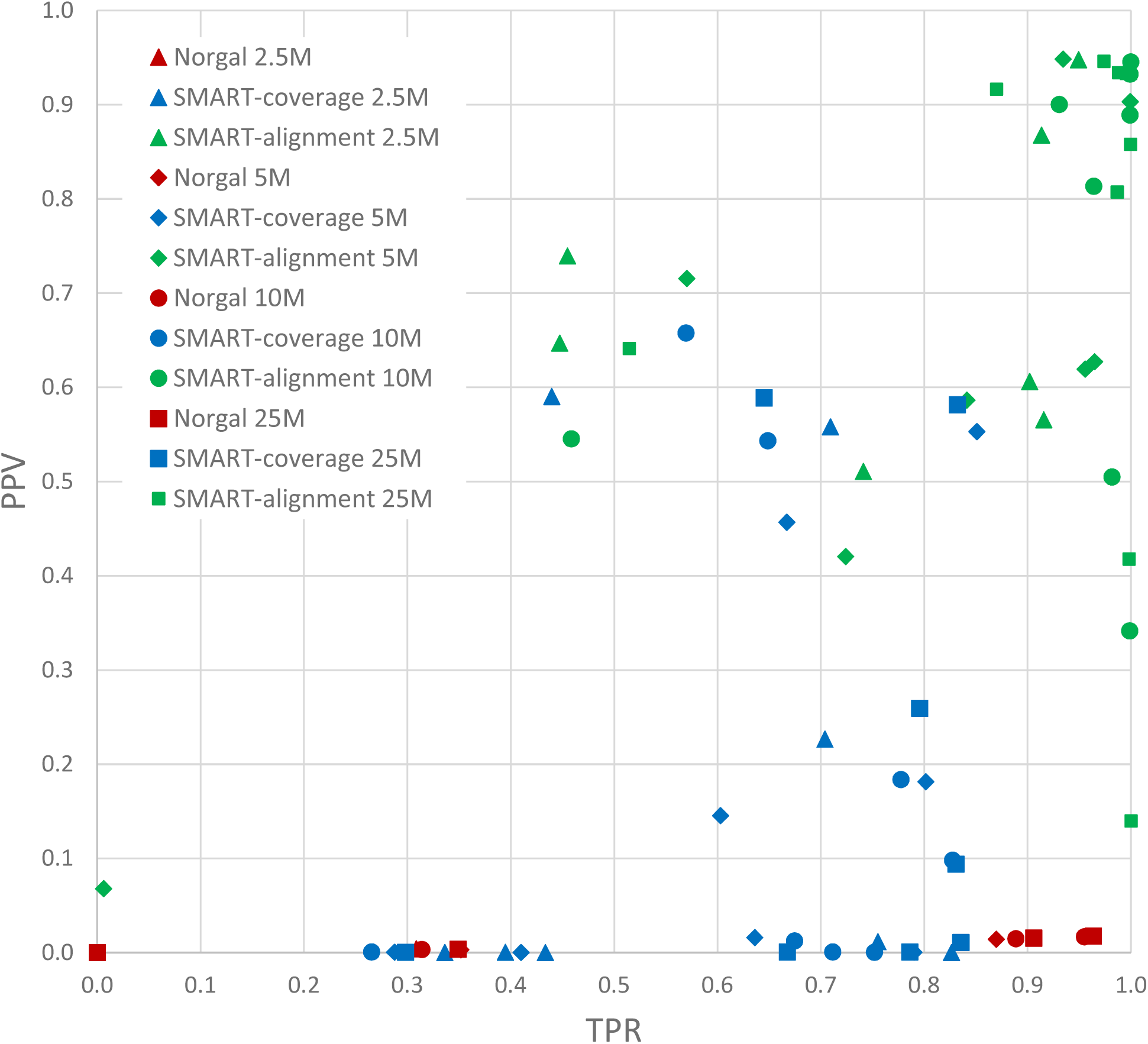
Assessment of read filtering accuracy for human datasets with 2.5-25M read pairs.

Compared to Norgal, the coverage-based filter of SMART performs much better at all sequencing depths. It has positive TPR on *all* datasets and at all sequencing depths, with the average TPR increasing from 0.575 ± 0.192 for 2.5M reads to 0.711 ± 0.183 for 25M reads. The filter also has better average PPV than Norgal, ranging from 0.173 ± 0.259 for 2.5M reads to 0.192 ± 0.258 for 25M reads, although the variability in PPV is quite high and many filtered reads sets still contain a majority of reads of nuclear origin. The coverage-based filter’s accuracy is likely to be negatively impacted by sequencing coverage non-uniformities of mitochondrial genome [37] as well as the presence in the nuclear genome of repeats with similar copy number. Both of these problems are mitigated by SMART’s alignment-based filter which has dramatically improved accuracy, with average TPR and PPV values ranging from 0.79 ± 0.222 and 0.728 ± 0.172, respectively, for 2.5M reads to 0.916 ± 0.168 and 0.708 ± 0.292 for 25M reads.

### Comparison with other tools

Table 3 reports the percentage identity, computed using Mauve [38], between the sequences reconstructed by each compared method on the human datasets described in Table 1 and the 1KGP ground truth. For each method and each dataset, the percentage identity is typeset in bold if the reconstructed sequence was a complete circular genome. Besides Norgal, NOVOPlasty, PlasmidSPAdes, and SMART we also ran MITObim in *de novo* mode but none of the MITObim runs completed successfully. The results show that when runs are successful, the quality of mitogenomes produced by Norgal, NOVOPlasty, PlasmidSPAdes, and SMART is very high. However, the success rates of different tools vary substantially. As mentioned above, MITObim *de novo* did not complete any of the 32 runs successfully. Norgal was successful in only 3 of the 32 runs (one with 10M read pairs and two with 25M pairs) but none of of the 3 assembled sequences was circular. Consistent to the very low read filtering PPV reported in Figure 3, in 19 of the runs Norgal generated nuclear contigs. NOVOplasty performed better, with 7 successful runs out of 32, and 6 of the 7 successful runs producing circular sequences. PlasmidSPAdes was successful in half of the runs, with 11 of the 16 successful runs producing circular mitogenomes. However, PlasmidSPAdes also had the highest running time, with four of the runs being stopped after 14 days. PlasmidSPAdes, which uses a negative read filtering strategy similar to Norgal’s, also generated nuclear contigs in a large number of runs (11 out of 32). SMART had the highest success rate on the human datasets, with 26 successful runs, of which all but one produced circular sequences. For all methods the success rate seems to increase with the sequencing depth, however SMART outperforms the other methods at each sequencing depth.

**Table 3:** Assembly accuracy comparison on human datasets. The percentage identity to the 1KGP reference, computed using Mauve [38] for each methods and each dataset, is typeset in bold when the reconstructed sequence is a complete circular genome.

| #Pairs | Run ID | Norgal | NOVOPlasty | PlasmidSPAdes | SMART |
| --- | --- | --- | --- | --- | --- |
| 2,500,000 | ERR020236 | - | - | **99.98** | **99.98** |
|  | SRR1596847 | nuclear | - | nuclear | - |
|  | ERR1044792 | nuclear | - | nuclear | **99.98** |
|  | SRR071189 | nuclear | - | 99.80 | **99.98** |
|  | ERR250974 | - | - | nuclear | - |
|  | ERR043002 | - | - | **99.96** | **99.95** |
|  | ERR047805 | nuclear | - | nuclear | - |
|  | ERR065367 | nuclear | - | nuclear | - |
| 5,000,000 | ERR020236 | - | 99.96 | **99.98** | **99.98** |
|  | SRR1596847 | nuclear | - | nuclear | 99.96 |
|  | ERR1044792 | nuclear | - | 99.98 | **99.98** |
|  | SRR071189 | nuclear | **99.96** | - | **99.98** |
|  | ERR250974 | - | - | nuclear | - |
|  | ERR043002 | - | - | **99.90** | **99.95** |
|  | ERR047805 | nuclear | - | nuclear | - |
|  | ERR065367 | nuclear | - | nuclear | **99.90** |
| 10,000,000 | ERR020236 | 99.98 | - | **99.98** | **99.98** |
|  | SRR1596847 | nuclear | - | **99.98** | **99.98** |
|  | ERR1044792 | nuclear | **99.97** | **99.98** | **99.98** |
|  | SRR071189 | nuclear | **99.96** | 99.97 | **99.98** |
|  | ERR250974 | - | - | 99.60 | **99.98** |
|  | ERR043002 | - | - | **99.95** | **99.90** |
|  | ERR047805 | nuclear | - | nuclear | **99.90** |
|  | ERR065367 | nuclear | - | timeout | **99.90** |
| 25,000,000 | ERR020236 | 99.98 | - | timeout | **99.98** |
|  | SRR1596847 | - | - | **99.98** | **99.97** |
|  | ERR1044792 | nuclear | **99.98** | **99.98** | **99.98** |
|  | SRR071189 | nuclear | **99.97** | 99.97 | **99.98** |
|  | ERR250974 | - | - | **99.90** | **99.98** |
|  | ERR043002 | 99.95 | **99.90** | timeout | **99.90** |
|  | ERR047805 | nuclear | - | nuclear | **99.90** |
|  | ERR065367 | nuclear | - | timeout | **99.90** |

### Seed effect

Requiring previous knowledge regarding the organism of interest in the form of a seed sequence is a drawback that SMART shares with seed-and-extend methods [7]. However, SMART uses the seed only to estimate the distribution of mtDNA *k*-mer counts and then extracts mtDNA reads based on their *k*-mer coverage instead of retrieving reads based on overlap with the seed sequence. We expect the SMART approach to work even with very short seed sequences such as the COI gene, and with seed sequences from other species. To assess the effect of seed sequence length and degree of dissimilarity, we ran SMART on 2.5M-25M read pairs randomly sampled from WGS sequencing run ERR020236 and using seed sequences of varying length and origin. Details on these seed sequences, including their lengths and accession numbers, are given in Table 4. In addition to four human COI gene sequences of 386-1542bp downloaded from NCBI GeneBank and the Barcode of Life Data System (BOLD) we included in the comparison a 386bp COI sequence from the 1KGP individual that was the source of the WGS sequencing data and six COI sequences from the four species most closely related to humans: three COI sequences from *Pan troglodytes*, and one sequence each from *Pan paniscus, Gorilla gorilla*, and *Gorilla beringei*.

**Table 4:**
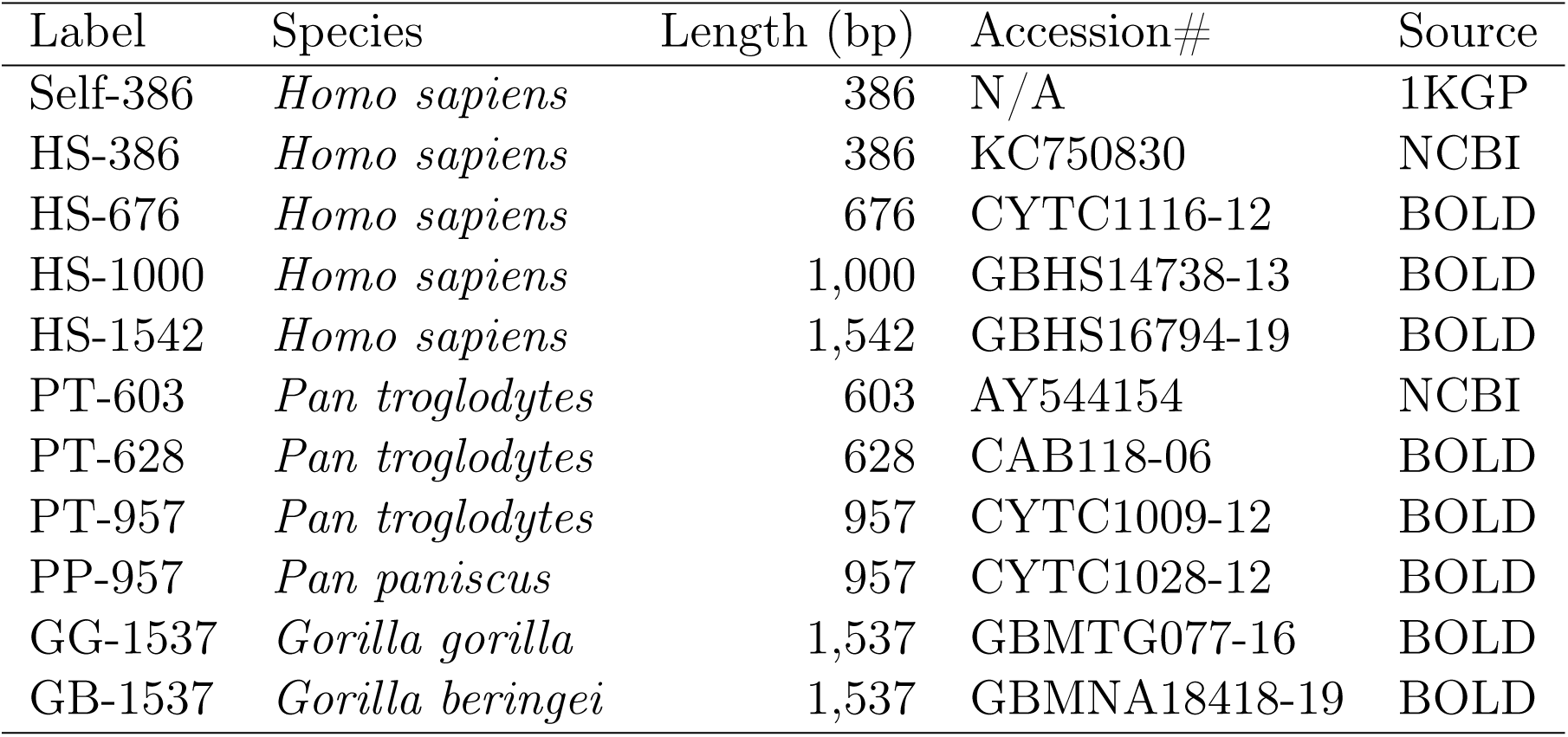
Seed sequences used to assess accuracy of SMART read filters on datasets with 2.5M-25M read pairs randomly selected from WGS run ERR020236.

| Label | Species | Length (bp) | Accession# | Source |
| --- | --- | --- | --- | --- |
| Self-386 | *Homo sapiens* | 386 | N/A | 1KGP |
| HS-386 | *Homo sapiens* | 386 | KC750830 | NCBI |
| HS-676 | *Homo sapiens* | 676 | CYTC1116-12 | BOLD |
| HS-1000 | *Homo sapiens* | 1,000 | GBHS14738-13 | BOLD |
| HS-1542 | *Homo sapiens* | 1,542 | GBHS16794-19 | BOLD |
| PT-603 | *Pan troglodytes* | 603 | AY544154 | NCBI |
| PT-628 | *Pan troglodytes* | 628 | CAB118-06 | BOLD |
| PT-957 | *Pan troglodytes* | 957 | CYTC1009-12 | BOLD |
| PP-957 | *Pan paniscus* | 957 | CYTC1028-12 | BOLD |
| GG-1537 | *Gorilla gorilla* | 1,537 | GBMTG077-16 | BOLD |
| GB-1537 | *Gorilla beringei* | 1,537 | GBMNA18418-19 | BOLD |

Figure 4 shows the number of mitochondrial (true positives, or TP) and nuclear (false positive, or FP) read pairs that pass the coverage- and alignment-based SMART filters, respectively. For all sequencing depths, the use of human seeds leads to recovery of almost all mitochondrial reads following the alignment based filter, with few FPs. For non-human seeds the low sensitivity of the coverage-based filter leads to more variable performance of the alignment-based filter although the number of FPs remains low. As shown in Figure 4 by the seed labels typeset in bold, SMART succeeded in assembling a complete circular mtDNA genome using all five human seeds, regardless of seed length and sequencing depth. SMART also has a less-consistent but still high success rate at assembling the complete circular mtDNA genome when using the seeds from related species. All reconstructed sequences had an overall average of 99.96% identity to the mitochondrial genome published by 1KGP for individual HG00501 as computed by Mauve [38]; the average percent identity is 99.98% for mitogenomes reconstructed using human seeds.

**Figure 4:**
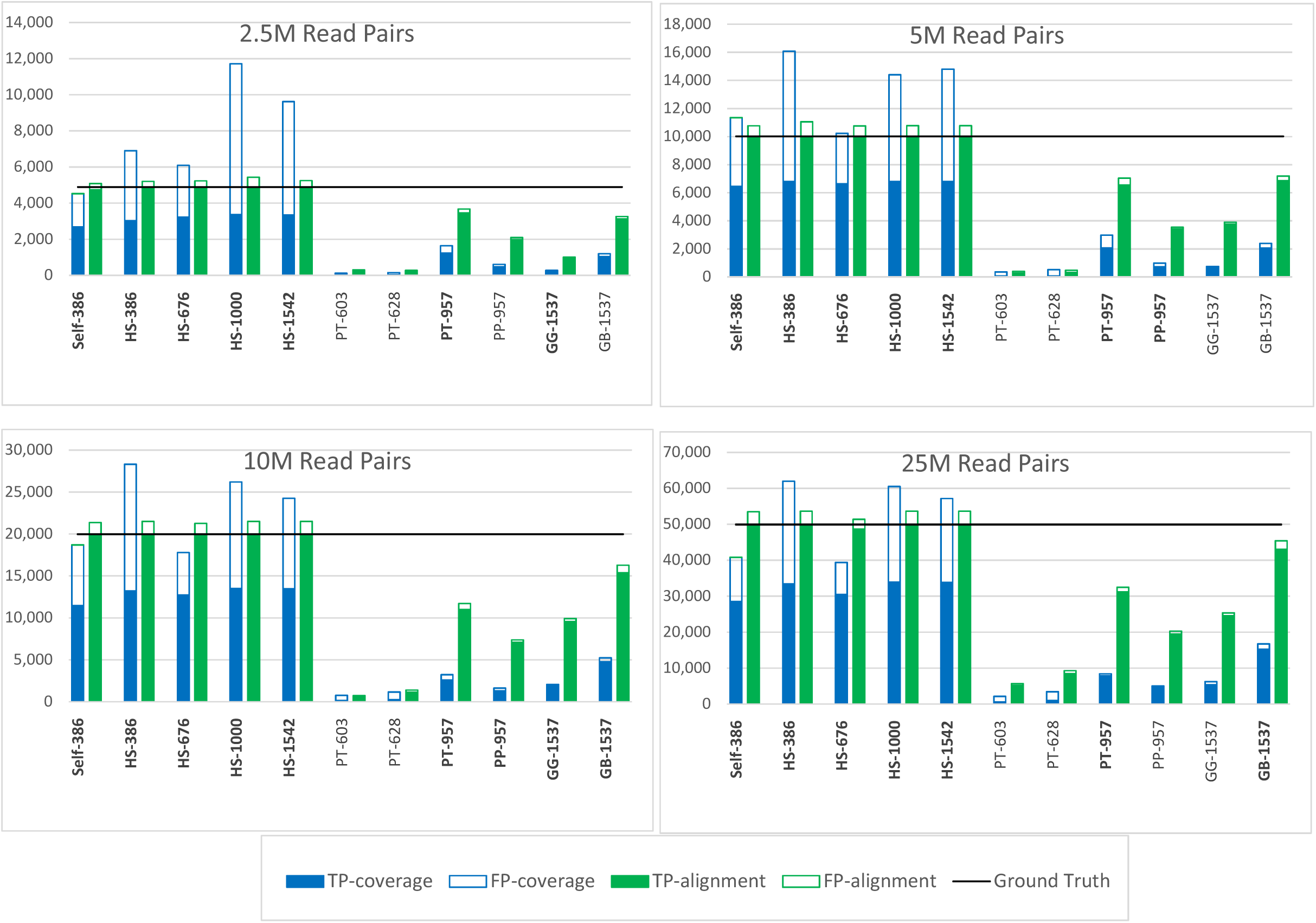
Effect of the seed on read filtering accuracy for 2.5M-25M read pairs randomly selected from WGS run ERR020236. X-axis represents the used seed sequences and y-axis represents the number of pair reads which are the output of coverage-based and reference-based read filter

### SMART results on other species

SMART retained high success rate and assembly accuracy when assembling mitogenomes for the 10 non-human datasets described in Table 2. All SMART assemblies except that of *Rana temporaria* were circular. Table 5 gives the percentage identity between the SMART assemblies and the NCBI mitogenomes of the corresponding species computed using five different tools: Mauve [38], LASTZ [39], MUSCLE [40], ClustalW [41], and MAFFT [32]. Although there are minor differences between the percentage identity reported by different tools, all values are very high, with the slightly lower identities observed for *Aspergillus niger, Grus japonensis*, and *Phlebotomus papatasi* possibly being explained by the higher intra-species variability within these species.

**Table 5:**
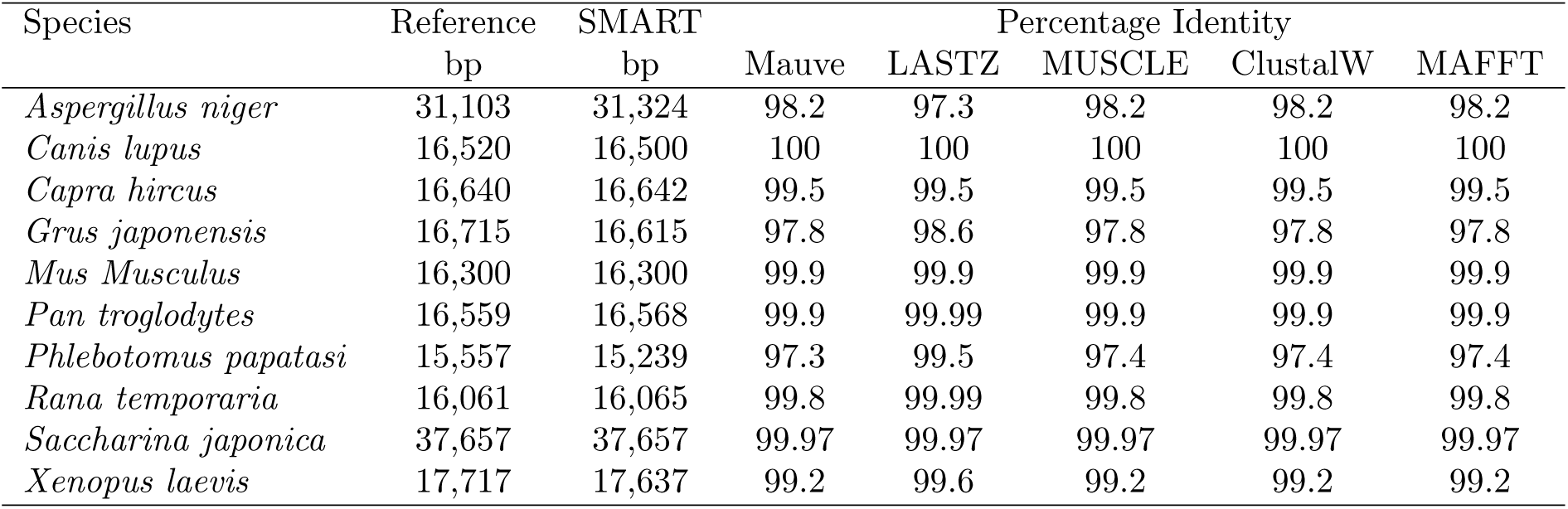
SMART assembly accuracy for non-human datasets. All SMART assemblies except *Rana temporaria* are circular.

### Conclusions

SMART is an automated pipeline for *de novo* mitogenome assembly from paired WGS reads. It is based on a novel statistical framework that includes probabilistic read filtering based on coverage, likelihood maximization for resolving ambiguities in the assembly graph, and assembly confidence estimation using bootstrapping. Experimental results on both human and non-human datasets show that SMART produces complete/circular assemblies with high success rate even for low-coverage WGS data and in presence of repeats. The pipeline is available via a user-friendly galaxy interface at https://neo.engr.uconn.edu/?toolid=SMART.

## Abbreviations

SMART: Statistical Mitogenomes Assembly with Repeats
1KGP: 1000 Genomes Project
ATP: Adenosine triphosphate
NGS: Next-Generation Sequencing
WGS: Whole Genome Sequencing
WES: Whole Exome Sequencing
COI: Cytochrome c oxidase I
NCBI: The National Center for Biotechnology Information
BLAST: Basic Local Alignment Search Tool
TPR: True Positive Rate
PPV: Positive Predictive Value

## Ethics approval and consent to participate

Not applicable.

## Consent to publish

Not applicable.

## Availability of data and material

Galaxy histories of the SMART runs for the ten non-human species are publicly accessible at https://neo.engr.uconn.edu/histories/list_published.

All data generated or analysed during this study are included in this published article.

## Author’s contributions

Both authors designed the SMART pipeline. FA implemented the pipeline and deployed it using the Galaxy framework. Both authors conducted the experiments, analyzed the data, and wrote the manuscript. Both authors have read and approved the final manuscript.

## Acknowledgements

A one-page abstract of this work was previously published in the Proceedings of the 8th IEEE International Conference on Computational Advances in Bio and Medical Sciences [42].

